# Bisphenol AF induces overactivation of primordial follicles via Hippo signaling and causes premature ovarian insufficiency in mice

**DOI:** 10.1101/2024.06.19.599666

**Authors:** Xiaoyang Liu, Mingxi Yu, Tiancheng Wang, Xiangdong Hu, Rui Zhong, Yuan Xiao, Yan Xu, Mei Zhang, Shuang Tang

## Abstract

Premature depletion of ovarian reserve is a cause of female infertility. Plasticizer Bisphenol AF (BPAF) residues have been found in human reproductive-related samples. However, little is known about its influence on the ovarian reserve. Here, we demonstrate that BPAF exposure causes excessive activation of primordial follicles through Hippo signaling in young females, resulting in rapid exhaustion of the ovarian reserve and, ultimately, premature ovarian insufficiency. We found that oral ingestion of BPAF disrupts normal estrous cyclicity and induces constant estrus in a dose-dependent manner. BPAF can upregulate the expression of YAP transcriptional coactivator. Intriguingly, only the high dose of BPAF inhibits Hippo signaling by eliciting a substantial decrease in phospho-YAP levels. Through such regulatory effects, BPAF causes YAP translocation into the nucleus and triggers the overactivation of primordial follicles. Collectively, this study proposes a novel toxicological mechanism explaining the negative impact of BPAF on the ovarian health of young females.

## Introduction

Ovulation defects are the leading cause of infertility in women, accounting for 30% of all cases of female infertility (1). The mammalian ovary accommodates dormant primordial follicles containing immature oocytes surrounded by granulosa cells. At reproductive age, a limited number of primordial follicles are recruited from the ovarian follicle reservoir during each estrus. Then, these follicles gradually grow into antral follicles in response to gonadotropins and ovulate mature oocytes ready for fertilization (2, 3). The initial activation of primordial follicles is strictly controlled by specific molecular cues, such as the Hippo or phosphatidylinositol 3-kinase (PI3K) pathways (4–7). This mechanism protects the female ovarian reserve, allowing for a relatively lengthy reproductive lifespan (2, 3). However, if the primordial follicles are stimulated by some external hazardous materials and then abnormally activated, it is likely to cause premature ovarian insufficiency (POI), also known as premature ovarian failure (POF) (8–11).

Plastic products are ubiquitous in modern life. Bisphenols are commonly used plasticizers that increase the flexibility and processability of plastics. Bisphenol A (BPA) was previously one of the most widely used plasticizers, which has yet been banned since 2010 due to its potential risks to human health (12, 13). Bisphenol AF (BPAF) is a fluorinated analog of BPA. It is applied in the manufacture of epoxy resins and polycarbonate polymers. The downstream products include, but are not limited to, water bottles, dental sealants, medical equipment, cosmetics, and reusable food containers (14–16). Consequently, BPAF residues have been detected in maternal plasma, urine, umbilical cord plasma, and placenta samples (17–19). It has been suggested that BPAF might be detrimental to ovarian health. For instance, BPAF is related to reduced oocyte quality due to its toxicity in inducing DNA damage, disturbing meiosis, and altering epigenetic modification (20–22). BPAF can also impair the viability of mammalian granulosa cells (23, 24). Moreover, a recent study showed that BPAF exposure affects ovarian morphology and alters mRNA expression profiles associated with ovarian diseases (25). However, little is known about the influence of BPAF on ovarian reserve. In this study, we investigated whether BPAF could affect the ovarian reserve of young adult females by using the mouse model. The results show that BPAF exposure triggered the overactivation of primordial follicles by upregulating the expression of YAP transcriptional coactivator and suppressing YAP phosphorylation in the ovary, ultimately leading to premature exhaustion of ovarian reserve and a substantial reduction in fecundity.

## Results

### BPAF exposure causes constant estrus in young adult mice

When investigating the impact of BPAF on embryonic development, an increased frequency of mating was observed in treated female mice. This led us to investigate the effects of BPAF on the estrous cyclicity. The estrous cycle was monitored by daily vaginal smearing and cytologic analyses over 30 days. Mice in the vehicle group displayed normal cycling through the four stages. Treatment with 10 mg/kg of BPAF (BPAF10) resulted in only mild interference with the estrous cycle progression, with changes not being significant. In contrast, exposure to 250 mg/kg of BPAF (BPAF250) induced constant estrus in treated females, beginning between the sixth and twelfth days of BPAF250 treatment (Figure 1A and Figure S1). Consistently, the number of estrous cycles decreased markedly over the 30-day treatment period (Figure 1B). These data manifest that BPAF exposure alters the estrous cyclicity in a dose-dependent manner. In the endocrine-disrupting chemical (EDC) control and exogenous estrogen control groups, treatment with BPA or estradiol disrupted estrous cycles and suppressed estrus in the treated mice (Figure 1A and Figure S1). It is noted that neither of these two groups exhibited the constant-estrus phenotype observed in the BPAF250 group. These results suggest that the constant estrus caused by BPAF exposure should not be solely attributed to the endocrine-disrupting effects of bisphenols, but rather may involve other regulatory mechanisms.

**Figure 1.**
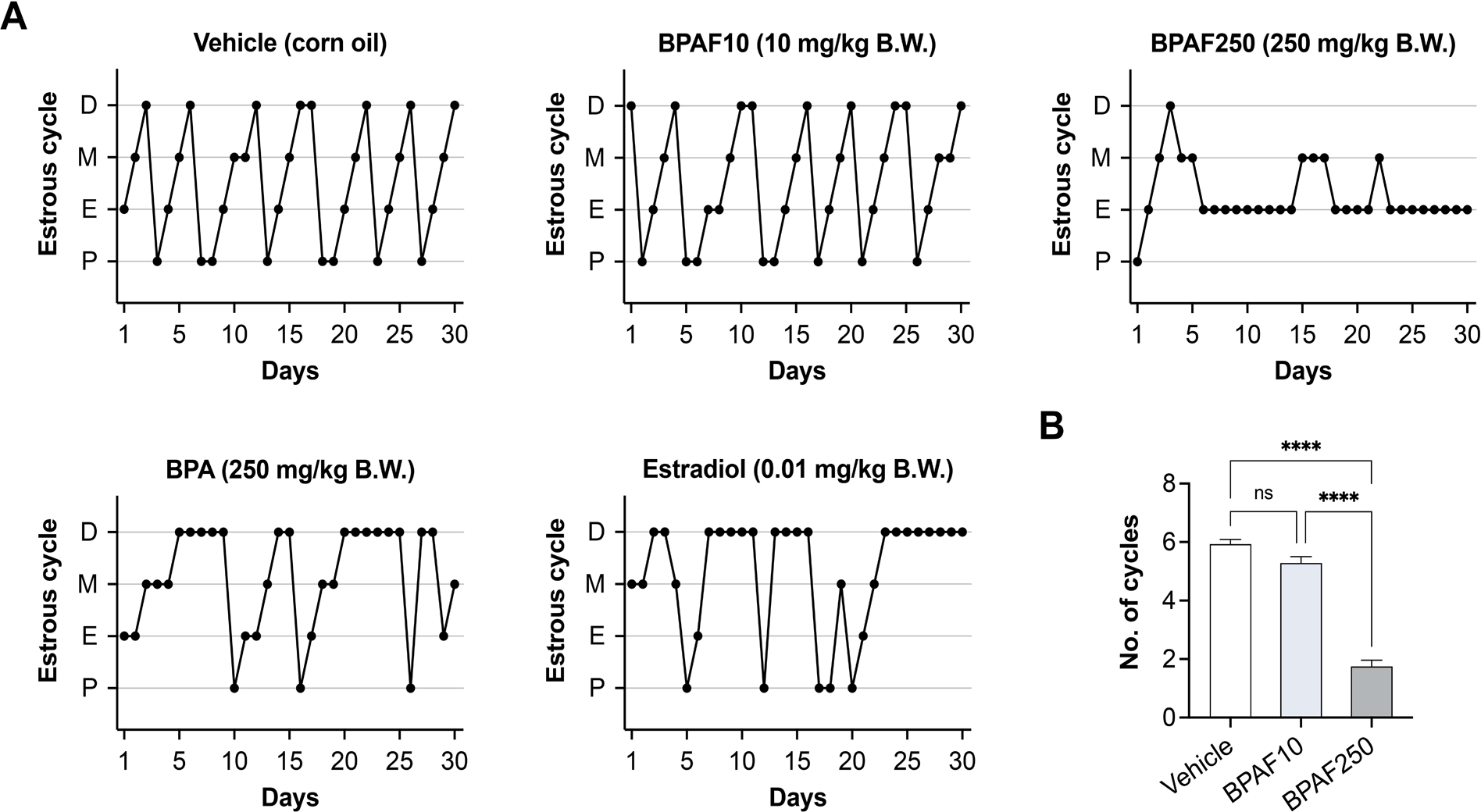
Exposure to BPAF causes aberrant constant estrus in young female mice. (A) Representative images illustrating estrous cyclicity patterns in mice (8-week-old) following intragastric administration of corn oil (Vehicle, n = 14), 10 mg/kg/day BPAF (BPAF10, n = 14), 250 mg/kg/day BPAF (BPAF250, n = 16), 250 mg/kg/day BPA (BPA, n = 9), or 0.01 mg/kg/day estradiol (Estradiol, n = 9) over a 30-day period. The stages of the estrous cycle are denoted as follows: P (proestrus), E (estrus), M (metestrus), and D (diestrus). The abbreviation B.W. refers to body weight. (B) The number of estrous cycles experienced by the treated mice during the 30-day treatment. The period from the initial estrus phase through diestrus and back to the subsequent estrus phase was counted as one cycle. The experiment was performed three times. Data were presented as mean ± SEM and analyzed by one-way ANOVA (Tukey’s multiple comparisons test). “ns” non-significant difference, **** *P* < 0.0001.

### BPAF exposure results in decreased fecundity and depleted ovarian reservoir in middle chronological age

Following 30 days of BPAF treatment, female mice underwent a 10-day recovery period. Subsequently, they were caged with males for a mating period of 12 weeks. The litter size and number of pups were recorded to assess the effect of BPAF on female fecundity. Although the mean size of the first litter was similar among the vehicle, BPAF10, and BPAF250 groups, there was a substantial reduction in the last litter size in the BPAF250 group (Figure 2A and B). Furthermore, the over-time record of the cumulative number of pups per female revealed a trend toward a declined fecundity in BPAF250 treated mice compared with females from the other two groups (Figure 2C). At the end of the 12-week surveillance, the mice were 6 months old, roughly comparable to 30-year-old human women in their middle ages (26). These data demonstrate that exposure to BPAF in young adulthood may result in decreased fecundity in middle chronological age.

**Figure 2.**
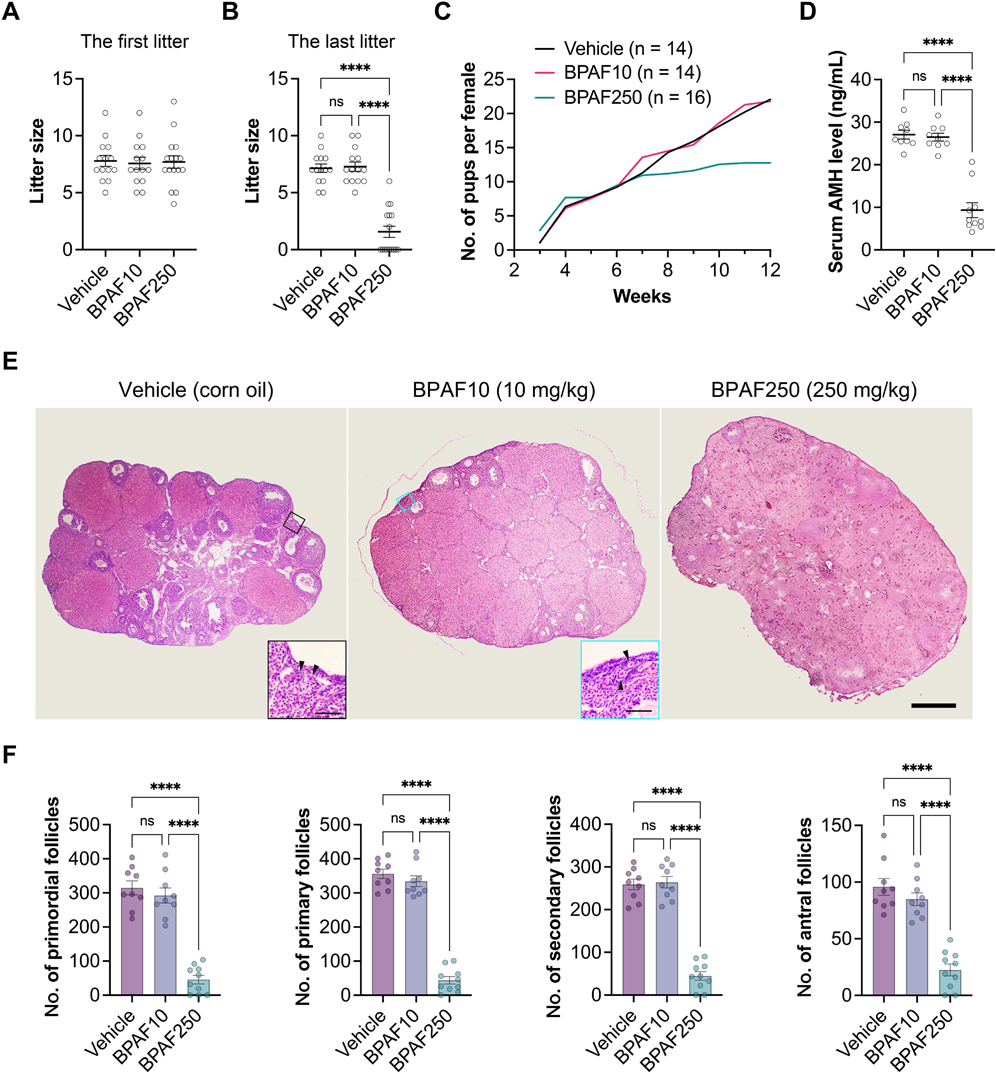
Exposure to BPAF results in decreased litter size and depleted ovarian reservoir in middle chronological age. Following 30 days of BPAF treatment, female mice underwent a 10-day recovery period and were then caged with males for a mating period of 12 weeks. (A) The first litter size of each group. Data were presented as mean ± SEM and analyzed by one-way ANOVA. No significant differences. (B) The last litter size of each group during the 12-week surveillance. (C) Comparison of the cumulative number of pups per female in each group. (D) Serum AMH levels in 6-month-old mice from each group at the end of the 12-week surveillance. (E) Representative micrographs of hematoxylin and eosin stained ovaries from 6-month-old mice in each group. Scale bar = 500 μm. The insets represent the enlargements of the boxed regions, showing the representative images of primordial follicles (arrowheads). Scale bar = 50 μm. (F) Numbers of different types of ovarian follicles. The experiments were repeated three times. Data were presented as mean ± SEM and analyzed by one-way ANOVA (Tukey’s multiple comparisons test). “ns” non-significant difference, **** *P* < 0.0001.

The ovaries of these mice were collected for histological examination. No significant changes in ovarian weight were observed among the three groups (Figure S2). To elucidate the cause underlying the decreased fecundity of BPAF250 mice, we compared the primordial, primary, secondary, and antral follicle numbers in ovarian sections from the three groups. In comparison with the vehicle and BPAF10 groups, numbers of different types of ovarian follicles were significantly decreased in the BPAF250 group (Figure 2E and F). Moreover, we measured serum anti-Mullerian hormone (AMH) levels as a marker of ovarian reserve exhaustion (27). The serum AMH level was markedly lower in BPAF250 treated mice in contrast to the mice with vehicle or BPAF10 treatment (Figure 2D). These findings indicate that exposure to BPAF in young adulthood results in ovarian reserve decline and, ultimately, POI in middle chronological age.

### BPAF triggers the overactivation and depletion of primordial follicles in young females

To further investigate the cause of ovarian reserve decline in middle-aged mice, we examined effects of BPAF on primordial follicle activation and follicular development in young adult mice. We had 8-week-old mice treated for 21 days and collected the ovaries for histological examination. The number of primordial follicles was significantly lower in the BPAF250 group compared with the vehicle and BPAF10 groups. In contrast, the primary, secondary, and antral follicle numbers in the BPAF250 group were markedly higher than those in the vehicle and BPAF10 groups (Figure 3A and B). No significant changes in ovarian weight were observed among the three groups (Figure S3). We further measured serum levels of follicle-stimulating hormone (FSH) and luteinizing hormone (LH) which are responsible for stimulating follicular growth and ovulation (28). In serum of BPAF250 treated mice, elevated levels of FSH and LH were observed relative to vehicle or BPAF10 treated mice (Figure 3C and D). Consequently, the follicle reservoir was quickly attrited under the BPAF challenge. These findings demonstrate that BPAF exposure triggers excessive activation of primordial follicles and promotes follicular development in young adult mice.

**Figure 3.**
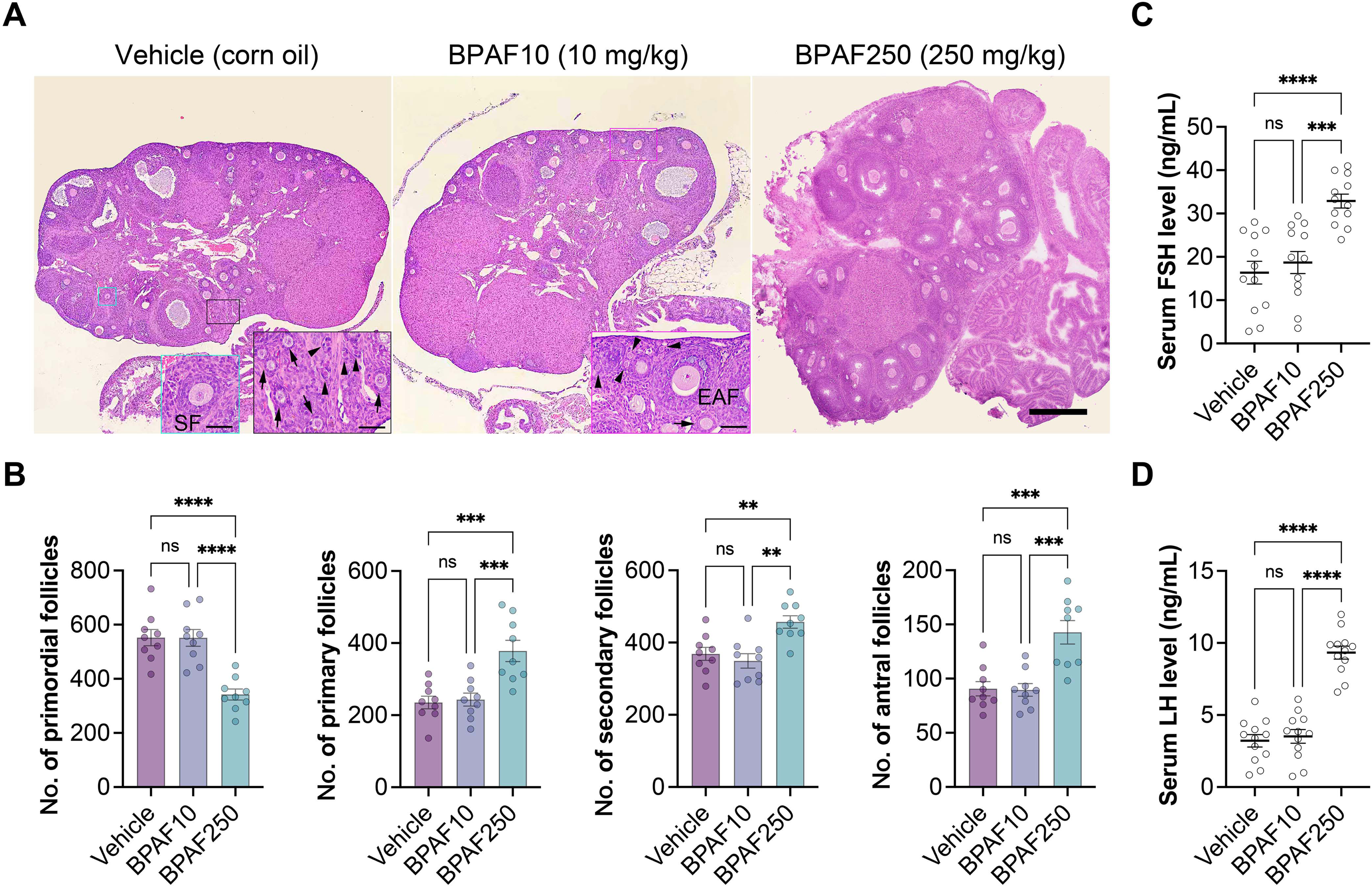
BPAF triggers excessive activation and depletion of primordial follicles in young females. (A) Representative micrographs of hematoxylin and eosin stained ovaries from mice (8-week-old) treated with corn oil (Vehicle, n = 9), 10 mg/kg/day BPAF (BPAF10, n = 9), or 250 mg/kg/day BPAF (BPAF250, n = 9) for 21 days. Scale bar = 500 μm. The insets represent the enlargements of the boxed regions. Arrowheads indicate primordial follicles. Arrows indicate primary follicles. SF, secondary follicle. EAF, early antral follicle. Scale bar = 50 μm. (B) Numbers of different types of follicles. (C and D) Serum FSH and LH levels in mice from the vehicle, BPAF10, and BPAF250 groups (n = 12 for each group). The experiments were repeated three times. Data were presented as mean ± SEM and analyzed by one-way ANOVA (Tukey’s multiple comparisons test). “ns” non-significant difference, ** *P* < 0.01, *** *P* < 0.001, **** *P* < 0.0001.

### BPAF upregulates YAP expression and suppresses YAP phosphorylation

In our recent work, we found that BPAF hinders trophectoderm differentiation in mouse preimplantation embryos through specific modulation of Hippo signaling. Therefore, we wanted to know whether BPAF-induced overactivation of primordial follicles is dependent on the Hippo signaling pathway. We assessed YAP expression and phosphorylation by immunoblot in ovaries isolated from the vehicle, BPAF10, and BPAF250 treated mice. Results demonstrated that ovarian YAP levels were significantly elevated in both BPAF10 and BPAF250 groups compared with the vehicle group (Figure 4A and B). In addition, qPCR analysis showed similar changes in *Yap* mRNA expression (Figure 4D). Notably, while YAP levels increased with escalating BPAF doses, the ascending rate plateaued, indicating that even low dose of BPAF is sufficient to induce YAP upregulation in the ovaries. There was no significant enhancement of this regulation with the increase of BPAF dosage (Figure 4B and D). Unlike YAP expression, the modulation of YAP phosphorylation state required high BPAF dosage. Specifically, relative to the vehicle group, there was no significant change in phospho-YAP (p-YAP) level in the BPAF10 group. However, a marked decrease in the level of p-YAP was observed in the BPAF250 group (Figure 4A and C). Thus, exposure to 250 mg/kg of BPAF leads to Hippo signaling off in the ovary. These results may explain why only BPAF250 treatment could induce constant estrus and primordial follicle activation (Figure 1 and Figure 3). Overall, BPAF possesses dual regulatory effects on the Hippo signaling pathway in a dose-related manner.

**Figure 4.**
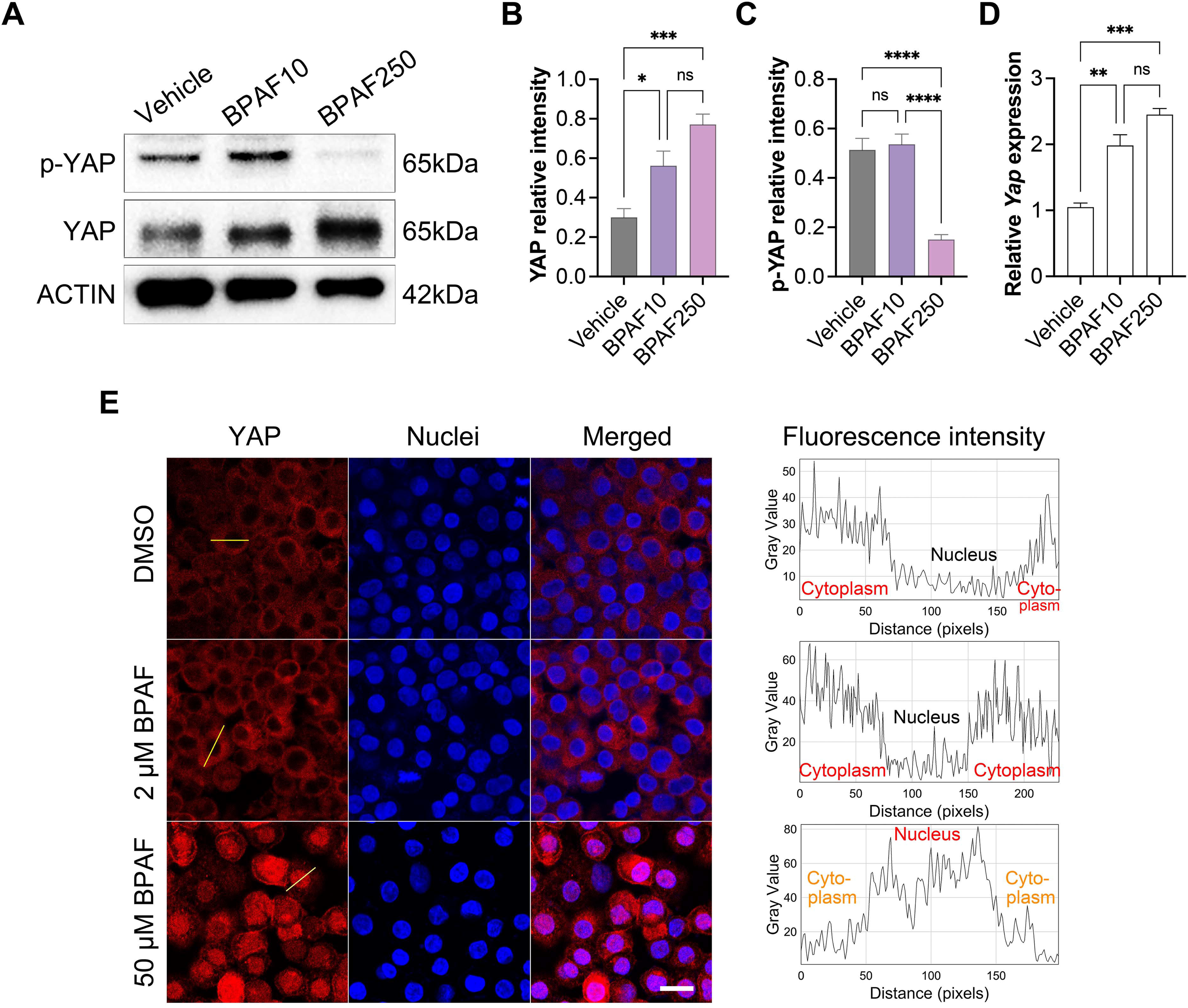
Effects of BPAF on YAP expression, phosphorylation, and cytoplasmic-nuclear translocation. (A) Representative immunoblot images of YAP and phospho-YAP (p-YAP) in ovaries from the vehicle, BPAF10, and BPAF250 groups. β-ACTIN was used as the loading control. (B and C) Densitometric analyses of protein expression levels of YAP and p-YAP normalized to ACTIN. (D) *Yap* mRNA levels in ovaries from each group were determined by qPCR analysis and the values were normalized to *Gapdh* expression. (E) The effect of BPAF on YAP localization in ovarian granulosa-like cells. Left: Representative confocal images of YAP distribution in cells exposed to DMSO, 2 μM, and 50 μM of BPAF. Scale bar = 20 μm. Right: Relative fluorescence intensity of YAP in the cytoplasm and the nucleus of cells exposed to different doses of BPAF. The fluorescence intensity of the line region across the entire cell (indicated by the yellow line in the left panel) was analyzed using the Plot Profile tool by ImageJ. DMSO, dimethyl sulfoxide. The experiments were repeated at least three times. Data for (B-D) were presented as mean ± SEM and analyzed by one-way ANOVA (Tukey’s multiple comparisons test). “ns” non-significant difference, * *P* < 0.05, ** *P* < 0.01, *** *P* < 0.001, **** *P* < 0.0001.

### BPAF exposure induces YAP nuclear translocation

In the ovary, suppression of Hippo signaling in squamous pre-granulosa cells induces YAP translocation into the nucleus to initiate the expression of downstream target genes, orchestrating profound changes necessary for follicle activation and development (4, 6, 7). Given this, we performed YAP immunostaining assays according to our initial design, yet the antibody did not work well on ovary sections. Therefore, for further analysis of YAP localization under the influence of BPAF, we instead performed YAP immunofluorescence on a human ovarian granulosa-like cell line COV434. As can be seen, in comparison to the cytoplasmic localization in DMSO and low dose (2uM BPAF) groups, Yap nuclear translocation occurred in cells from the high dose (50uM BPAF) group (Figure 4E). These findings can be mutually corroborated with the above results and demonstrate that BPAF affects YAP cytoplasmic-nuclear translocation in a dose-related manner. Another important implication of these findings is that the mechanism we revealed here might also be conserved in humans. In summary, BPAF inhibits the Hippo signaling pathway in the ovary through upregulation of YAP expression, downregulation of YAP phosphorylation, and induction of YAP nuclear translocation.

## Discussion

Premature exhaustion of ovarian follicle reservoir is a major culprit of female infertility. Environmental pollutants may affect the declining trajectory of ovarian reserve, suggesting a possible etiology for POI (9–11). Because the plasticizer BPAF is widely used, its residue is present in a variety of samples from reproductive-aged women (17–19). Therefore, it is imperative to investigate its influence on ovarian health. This study found that BPAF, in a dose-dependent manner, disrupted the normal estrous cyclicity and caused constant estrus, resulting in premature exhaustion of the ovarian reserve and a substantial reduction in fecundity. The dual regulatory effects of BPAF on the Hippo signaling play key roles during this process. BPAF can upregulate the expression of YAP transcriptional coactivator in the ovaries of treated mice. Intriguingly, only the high dose of BPAF inhibits Hippo signaling by eliciting a substantial decrease in phospho-YAP (p-YAP) levels. Through such regulatory effects, BPAF causes YAP translocation into the nucleus and triggers the overactivation of primordial follicles. These findings, to our knowledge, constitute the first report that revealed a close relevance of BPAF exposure to POI and unraveled the toxicological mechanism underlying the unfavorable impact of BPAF residues on the ovarian reservoir of young females.

## Materials and Methods

### Reagents

All chemicals were purchased from Sigma-Aldrich (St. Louis, MO) unless otherwise stated. Fetal bovine serum (FBS) and DMEM-HG medium were purchased from Gibco (Thermo Fisher Scientific, Waltham, MA). BPAF (CAS 1478-61-1, Lot: E1727018, Cat No. H106097, purity > 98.0%), BPA (CAS 80-05-7, Lot: L1803091, Cat No. B108652, purity > 99.0%) were purchased from Aladdin Scientific (Shanghai, China). β-Estradiol (CAS 50-28-2, Lot: CIPDG-PQ, Cat No. E0025, purity > 97.0%) were purchased from Tokyo Chemical Industry (Shanghai) Ltd. (Shanghai, China).

### Animal husbandry

CD-1 mice, 6-week-old females and 12-week-old males, were purchased from Beijing Vital River Laboratory Animal Technology Co., Ltd. (Beijing, China). The mice were kept on a 12-h light/dark cycle in a temperature-controlled room (23–25°C) of the SPF (specific pathogen-free) Lab Animal Center of the university. Animals were randomly group-housed four to a cage with food and water available *ad libitum* in bisphenol-free conditions. All experiments were conducted following the Guide for the Care and Use of Laboratory Animals and approved by the Institutional Animal Care and Use Committee of Shenyang Agricultural University (approval no. 12003).

### BPAF treatment regimen

The BPAF toxicity data for mice are currently limited. The LD50 (median lethal dose) of BPAF in rats following oral exposure is 3400 mg/kg body weight (mg/kg B.W. or abbreviated as mg/kg) (29), while the relevant dosage for mice remains unreported. In the present study, we conducted a preliminary experiment in mice and found that the dose of 2500 mg/kg resulted in lethality in about half of the mice under our laboratory conditions. Consequently, one-tenth of this dose, namely 250 mg/kg, was used as the high dosage. In previous studies, BPAF was commonly administered at doses of 2–300 mg/kg//day to test its reproductive toxicity (30). When abnormal estrous cycles and reduced pregnancy rates were observed in female rats, the no-observed-adverse-effect-level (NOAEL) was designated at the dose of 10 mg/kg/day or < 30 mg/kg/day (30). Referring to this, we set 10 mg/kg/day as the low dosage. Given the endocrine-disrupting effects of bisphenols, endocrine-disrupting chemical (EDC) control and exogenous estrogen control groups were established using BPA and estradiol, respectively. The oral LD50 of BPA is 2500 mg/kg for the mice (31), so one-tenth of the dose, 250 mg/kg, was utilized. The estradiol dosage was determined by reference to human clinical dosage (0.005–0.016 mg/kg/day), with adjustments made for equivalent dose of the drug based on body surface area ratio (9.1 times), resulting in an average dose of approximate 0.1 mg/kg/day. One-tenth of this dose, namely 0.01 mg/kg/day, was utilized.

In experiment 1, 8-week-old female mice were randomly assigned to five groups: estradiol control group (0.01 mg/kg/day, n = 3), BPA control group (250 mg/kg/day, n = 3), corn oil vehicle control group (n = 4–6), low-dose BPAF experimental group (10 mg/kg/day, n = 4–6), and high-dose BPAF experimental group (250 mg/kg/day, n = 4–6). The mice received intragastric administration daily from 1630 to 1730 h for 30 days, with the experiment being repeated three times.

In experiment 2, 8-week-old female mice were randomly assigned to three groups: corn oil vehicle control group (n = 4), low-dose BPAF experimental group (10 mg/kg/day, n = 4), and high-dose BPAF experimental group (250 mg/kg/day, n = 4). The mice received intragastric administration daily from 1630 to 1730 h for 21 days, with the experiment being repeated six times.

### Estrous cycle identification by vaginal cytology

A pipette was used to absorb 30 μL of normal saline, which was then pipetted several times at the vaginal opening to collect the shed cells. These collected cells were then placed on a slide and examined under a Nikon Ni-U upright microscope to observe their morphology and determine the specific estrous phase. The proestrus (P) has nucleated epithelial cells with no or few cornified cells; the estrus (E) mainly contains anucleated cornified cells; the metestrus (M) contains nucleated epithelial cells, anucleated cornified cells, and leukocytes; and the diestrus (D) mainly has leukocytes. The estrous cycle was monitored daily from 0800 to 0900 h over 30 days. Vaginal cytology was read and confirmed by at least two people blinded to the animal treatment.

### Monitoring of litter size

Following 30 days of intragastric administration, the mice underwent a 10-day recovery period equivalent to two estrous cycles. Then, the fecundity of these females was assessed through mating with a male in a ratio of 3: 1. The pregnant females were maintained individually in cages. Upon parturition, the pups were checked for the total number and removed. Meanwhile, the females were immediately returned to the breeding cage, allowing them to continue mating with the male. Over a 12-week observation period, the litter sizes of each female were recorded. If a female did not conceive or give birth during the last four weeks, the size of its last litter was assumed to be zero.

### Measurement of serum hormone levels

Before sacrifice, blood was collected from each anesthetized mouse. After coagulation, whole blood was centrifuged at 1,000*g* for 10 min. The supernatant serum was stored at −80°C until analysis. The measurement of AMH, FSH, or LH levels was conducted with the Mouse Anti-Mullerian Hormone ELISA Kit (E-EL-M3015), Mouse Follicle Stimulating Hormone ELISA Kit (E-EL-M0511), or Mouse Luteinizing Hormone ELISA Kit (E-EL-M3053) according to the manufacturer’s protocols (Elabscience, Houston, TX).

### Histology

After sacrifice, ovaries were harvested and weighed. Some ovaries were fixed in 10% neutral buffered formalin, dehydrated through ethanol gradients, embedded in paraffin, and sliced into 8-μm sections. Ovary sections were stained with hematoxylin and eosin (H&E). The number of primordial follicles was assessed in every two serial sections. The number of primary follicles, secondary follicles, and antral follicles was assessed in every five serial sections. Researchers who were unaware of the animal status and treatment counted ovarian follicles.

### Western blotting (WB)

The total proteins from the ovaries were separated by 10% sodium dodecyl sulfate polyacrylamide gel electrophoresis and transferred onto a polyvinylidene fluoride membrane (Millipore, Bedford, MA). The membrane was blocked in TBST buffer (20 mM Tris-HCl, pH 8.2, 150 mM NaCl, 0.1% Tween-20) containing 5% (w/v) non-fat dried milk for 1 h. Then, the membrane was incubated with primary antibodies in TBST containing 5% milk overnight at 4°C. The primary antibodies were mouse anti-YAP (H00010413-M01, RRID: AB_535096, 1: 1,500 dilutions) from Abnova; rabbit anti-phospho-YAP (4911, RRID: AB_2218913, 1: 10,000 dilutions) from Cell Signaling Technology; moues anti-β-ACTIN (60008-1, RRID: AB_2289225, 1: 20,000 dilutions) from Proteintech. After thorough washes in TBST, the membrane was incubated with horseradish peroxide-conjugated secondary antibodies (Beyotime, Jiangsu, China) at 1: 2,000 dilutions in TBST containing 5% milk for 1 h. After thorough washes in TBST, the membrane was developed using the SuperSignal^TM^ West Pico Substrate (Pierce, Rockford, IL). The densitometric quantification was determined by using ImageJ (1.53t) software (https://imagej.nih.gov/ij/), calculating the ratio of the protein to ACTIN intensity.

### Real-time quantitative RT-PCR (qPCR)

Total RNA from the ovaries was extracted using Trizol (Thermo Fisher). Reverse transcription was performed using the GoScrip Reversion Transcription System (Promega, Madison, WI). The qPCR was performed using SYBR Premix Ex Taq (Takara, Osaka, Japan). The reaction was carried out on the CFX96 Real-time Detection System (Bio-Rad, Hercules, CA) with the following primers: *Gapdh*: forward (F), 5’-GTGTTCCTACCCCCAATGTGT-3’ and reverse (R), 5’-ATTGTCATACCAGGAAATGAGCTT-3’; *Yap*: F, 5’-GGACCCTCGTTTTGCCATGAA-3’ and R, 5’-GCAGAGCTAATTCCTGTGGTC-3’. The program for quantitative PCR was as follows: 95°C × 30 s, followed by 40 cycles of 95°C × 5 s, 60°C × 30 s. Relative changes in mRNA expression were calculated by the 2^−ΔΔCt^ method using *Gapdh* as the reference gene.

### Cell culture and immunofluorescence (IF)

The human ovarian granulosa-like tumor cell line COV434 (RRID: CVCL_2010) was supplied by courtesy of Rosetta Stone Biotechnology Co., Ltd. (Taiyuan, China). The cells were cultured in 35 mm dishes with a 14 mm bottom well in DMEM-HG containing 10% FBS at 37°C under a humidified atmosphere with 5% CO_2_. When the density reached 70–80%, the cells were treated with 0, 2.5, and 50 μM of BPAF for 16 h. The immunofluorescence was performed with the routine method. For the negative control, all steps are the same except for the omission of the primary antibody. The cells were fixed in 4% paraformaldehyde in PBS for 20 min and permeabilized in 0.2% Triton X-100 in PBS for 5 min. Then, the cells were blocked in 0.3% bovine serum albumin (BSA) in PBS (blocking solution) for 1 h at room temperature. Without wash, the cells were incubated with mouse anti-YAP primary antibody (1: 400 dilutions, H00010413-M01, RRID: AB_535096, Abnova) in blocking solution overnight at 4°C. After thorough washes in PBS, the cells were incubated with Cy3-conjugated donkey anti-mouse secondary antibodies (1: 800 dilutions, 715-165-151, RRID: AB_2315777, Jackson ImmunoResearch) for 20 min in the dark. The nuclei were counterstained with 10 µg/ml DAPI (4’,6-diamidino-2-phenylindole) in PBS containing 0.5 mg/mL of DNase-free RNase A for 20 min in the dark. The cells were observed under a Nikon A1^+^ laser scanning confocal microscope (Nikon, Tokyo, Japan). For quantitative analysis, the fluorescence images were analyzed using the Plot Profile tool by ImageJ (1.53t) software (https://imagej.nih.gov/ij/).

### Statistical analysis

Each experiment was independently performed at least three times. The data are presented as mean ± standard error of mean (SEM). Multiple comparisons were analyzed by one-way ANOVA (Tukey’s multiple comparisons test) using GraphPad Prism 9 (GraphPad Software, La Jolla, CA). Values of *P* < 0.05 were considered significantly different. Statistically values of non-significant difference, *P* < 0.05, *P* < 0.01, *P* < 0.001, and *P* < 0.0001 were indicated by (ns), asterisks (*), (**), (***), and (****), respectively.

## Acknowledgment

We would like to thank Dr. Haifeng Li for his assistance in ovarian histology. We are grateful to Ms. Lili Ren for her work in language editing. This work was supported by the Liaoning Science Planning Project from the Educational Department of Liaoning Province (LSNJC202009) and the Natural Science Fund of the College of Bioscience and Biotechnology SYAU.

---

**Figure S1. Estrous cyclicity patterns in treated mice.** One repeated experiment was depicted as a representative. Female mice (8-week-old) were orally administered with corn oil (vehicle control, n = 6), 10 mg/kg/day BPAF (low-dose BPAF treatment, n = 6), 250 mg/kg/day BPAF (high-dose BPAF treatment, n = 6), 250 mg/kg/day BPA (endocrine-disrupting chemical control, n = 3), or 0.01 mg/kg/day estradiol (exogenous estrogen control, n = 3) for a duration of 30 days. P, proestrus; E, estrus; M, metestrus; D, diestrus. B.W., body weight.

**Figure S2. Comparison of the ovarian weight in middle chronological age.** (A) The ovary weight in the vehicle, BPAF10, and BPAF250 groups at the end of the 12-week surveillance. (B) Ovary weight normalized to body weight (B.W.) of mice from the three groups. Data were presented as mean ± SEM and analyzed by one-way ANOVA (Tukey’s multiple comparisons test). No significant differences.

**Figure S3. The ovarian weight in young adult mice exposed to BPAF.** (A) The ovary weight in mice with vehicle, BPAF10, and BPAF250 treatment for 21 days. (B) Ovary weight normalized to body weight (B.W.) of mice from the three groups. Data were presented as mean ± SEM and analyzed by one-way ANOVA (Tukey’s multiple comparisons test). No significant differences.

## References

1. Land, K. L., Miller, F. G., Fugate, A. C., Hannon, P. R., The effects of endocrine-disrupting chemicals on ovarian- and ovulation-related fertility outcomes. Molecular Reproduction and Development, 2022, 89(12), 608–631. doi: 10.1002/mrd.23652.

2. Telfer, E. E., Grosbois, J., Odey, Y. L., Rosario, R., Anderson, R. A., Making a good egg: human oocyte health, aging, and in vitro development. Physiological Reviews, 2023, 103(4), 2623–2677. doi: 10.1152/physrev.00032.2022.

3. Monniaux, D., Clément, F., Dalbiès-Tran, R., Estienne, A., Fabre, S., Mansanet, C., Monget, P., The Ovarian Reserve of Primordial Follicles and the Dynamic Reserve of Antral Growing Follicles: What Is the Link?1. Biology of Reproduction, 2014, 90(4). doi: 10.1095/biolreprod.113.117077.

4. Hsueh, A. J. W., Kawamura, K., Hippo signaling disruption and ovarian follicle activation in infertile patients. Fertility and Sterility, 2020, 114(3), 458–464. doi: 10.1016/j.fertnstert.2020.07.031.

5. Reddy, P., Liu, L., Adhikari, D., Jagarlamudi, K., Rajareddy, S., Shen, Y., Du, C., Tang, W., Hämäläinen, T., Peng, S. L., Lan, Z.-J., Cooney, A. J., Huhtaniemi, I., Liu, K., Oocyte-Specific Deletion of Pten Causes Premature Activation of the Primordial Follicle Pool. Science, 2008, 319(5863), 611–613. doi: 10.1126/science.1152257.

6. De Roo, C., Lierman, S., Tilleman, K., De Sutter, P., In-vitro fragmentation of ovarian tissue activates primordial follicles through the Hippo pathway. Human Reproduction Open, 2020, 2020(4), 1–16. doi: 10.1093/hropen/hoaa048.

7. Grosbois, J., Devos, M., Demeestere, I., Implications of Nonphysiological Ovarian Primordial Follicle Activation for Fertility Preservation. Endocrine Reviews, 2020, 41(6), 847–872. doi: 10.1210/endrev/bnaa020.

8. Béranger, R., Hoffmann, P., Christin-Maitre, S., Bonneterre, V., Occupational exposures to chemicals as a possible etiology in premature ovarian failure: A critical analysis of the literature. Reproductive Toxicology, 2012, 33(3), 269–279. doi: 10.1016/j.reprotox.2012.01.002.

9. Vabre, P., Gatimel, N., Moreau, J., Gayrard, V., Picard-Hagen, N., Parinaud, J., Leandri, R. D., Environmental pollutants, a possible etiology for premature ovarian insufficiency: a narrative review of animal and human data. Environmental Health, 2017, 16(1), 37. doi: 10.1186/s12940-017-0242-4.

10. Zhu, X., Liu, M., Dong, R., Gao, L., Hu, J., Zhang, X., Wu, X., Fan, B., Chen, C., Xu, W., Mechanism Exploration of Environmental Pollutants on Premature Ovarian Insufficiency: a Systematic Review and Meta-analysis. Reproductive Sciences, 2023, 31(1), 99–106. doi: 10.1007/s43032-023-01326-5.

11. Richardson, M. C., Guo, M., Fauser, B. C. J. M., Macklon, N. S., Environmental and developmental origins of ovarian reserve. Human Reproduction Update, 2013, 20(3), 353–369. doi: 10.1093/humupd/dmt057.

12. Ao, J., Huo, X., Zhang, J., Mao, Y., Li, G., Ye, J., Shi, Y., Jin, F., Bao, S., Zhang, J., Environmental exposure to bisphenol analogues and unexplained recurrent miscarriage: A case-control study. Environmental Research, 2022, 204, 112293. doi: 10.1016/j.envres.2021.112293.

13. Siracusa, J. S., Yin, L., Measel, E., Liang, S., Yu, X., Effects of Bisphenol A and its Analogs on Reproductive Health: A Mini Review. Reproductive Toxicology, 2018, 79, 96–123. doi: 10.1016/j.reprotox.2018.06.005.

14. Hercog, K., Maisanaba, S., Filipič, M., Sollner-Dolenc, M., Kač, L., Žegura, B., Genotoxic activity of bisphenol A and its analogues bisphenol S, bisphenol F and bisphenol AF and their mixtures in human hepatocellular carcinoma (HepG2) cells. Science of the Total Environment, 2019, 687, 267–276. doi: 10.1016/j.scitotenv.2019.05.486.

15. Bala, R., Singh, V., Rajender, S., Singh, K., Environment, Lifestyle, and Female Infertility. Reproductive Sciences, 2021, 28(3), 617–638. doi: 10.1007/s43032-020-00279-3.

16. Lehmler, H.-J., Liu, B., Gadogbe, M., Bao, W., Exposure to Bisphenol A, Bisphenol F, and Bisphenol S in U.S. Adults and Children: The National Health and Nutrition Examination Survey 2013–2014. ACS Omega, 2018, 3(6), 6523–6532. doi: 10.1021/acsomega.8b00824.

17. Pan, Y., Deng, M., Li, J., Du, B., Lan, S., Liang, X., Zeng, L., Occurrence and Maternal Transfer of Multiple Bisphenols, Including an Emerging Derivative with Unexpectedly High Concentrations, in the Human Maternal–Fetal–Placental Unit. Environmental Science & Technology, 2020, 54(6), 3476–3486. doi: 10.1021/acs.est.0c00206.

18. Chen, D., Kannan, K., Tan, H., Zheng, Z., Feng, Y.-L., Wu, Y., Widelka, M., Bisphenol Analogues Other Than BPA: Environmental Occurrence, Human Exposure, and Toxicity—A Review. Environmental Science & Technology, 2016, 50(11), 5438–5453. doi: 10.1021/acs.est.5b05387.

19. Zhang, B., He, Y., Zhu, H., Huang, X., Bai, X., Kannan, K., Zhang, T., Concentrations of bisphenol A and its alternatives in paired maternal-fetal urine, serum and amniotic fluid from an e-waste dismantling area in China. Environment International, 2020, 136, 105407. doi: 10.1016/j.envint.2019.105407.

20. Abdallah, S., Jampy, A., Moison, D., Wieckowski, M., Messiaen, S., Martini, E., Campalans, A., Radicella, J. P., Rouiller-Fabre, V., Livera, G., Guerquin, M.-J., Foetal exposure to the bisphenols BADGE and BPAF impairs meiosis through DNA oxidation in mouse ovaries. Environmental Pollution, 2023, 317, 120791. doi: 10.1016/j.envpol.2022.120791.

21. Ding, Z.-M., Jiao, X.-F., Wu, D., Zhang, J.-Y., Chen, F., Wang, Y.-S., Huang, C.-J., Zhang, S.-X., Li, X., Huo, L.-J., Bisphenol AF negatively affects oocyte maturation of mouse in vitro through increasing oxidative stress and DNA damage. Chemico-Biological Interactions, 2017, 278, 222–229. doi: 10.1016/j.cbi.2017.10.030.

22. Hiradate, Y., Hoshino, Y., Kobayashi, N., Nakano, K., Nishio, M., Sato, E., Tanemura, K., Comparison of the effects of BPA and BPAF on oocyte spindle assembly and polar body release in mice. Zygote, 2016, 24(2), 172–180. doi: 10.1017/S0967199415000027.

23. Huang, M., Li, X., Jia, S., Liu, S., Fu, L., Jiang, X., Yang, M., Bisphenol AF induces apoptosis via estrogen receptor beta (ERβ) and ROS-ASK1-JNK MAPK pathway in human granulosa cell line KGN. Environmental Pollution, 2021, 270, 116051. doi: 10.1016/j.envpol.2020.116051.

24. Bujnakova Mlynarcikova, A., Scsukova, S., Bisphenol analogs AF and S: Effects on cell status and production of angiogenesis-related factors by COV434 human granulosa cell line. Toxicology and Applied Pharmacology, 2021, 426, 115634. doi: 10.1016/j.taap.2021.115634.

25. Yue, H., Yang, X., Wu, X., Tian, Y., Xu, P., Sang, N., Identification of risk for ovarian disease enhanced by BPB or BPAF exposure. Environmental Pollution, 2023, 319, 120980. doi: 10.1016/j.envpol.2022.120980.

26. Flurkey, K. M., Currer, J., Harrison, D. E., “Chapter 20 - Mouse Models in Aging Research” in The Mouse in Biomedical Research (Second Edition), J. G. Fox, M. T. Davisson, F. W. Quimby, S. W. Barthold, C. E. Newcomer, A. L. Smith, Eds. (Academic Press, Burlington, 2007), 10.1016/B978-012369454-6/50074-1, pp. 637–672.

27. van Rooij, I. A., Broekmans, F. J., te Velde, E. R., Fauser, B. C., Bancsi, L. F., de Jong, F. H., Themmen, A. P., Serum anti-Müllerian hormone levels: a novel measure of ovarian reserve. Human Reproduction, 2002, 17(12), 3065–3071. doi: 10.1093/humrep/17.12.3065.

28. Hillier, S. G., Gonadotropic control of ovarian follicular growth and development. Molecular and Cellular Endocrinology, 2001, 179(1), 39–46. doi: 10.1016/S0303-7207(01)00469-5.

29. Waidyanatha, S., Collins, B. J., Cunny, H., Aillon, K., Riordan, F., Turner, K., McBride, S., Betz, L., Sutherland, V., An investigation of systemic exposure to bisphenol AF during critical periods of development in the rat. Toxicology and Applied Pharmacology, 2021, 411, 115369. doi: 10.1016/j.taap.2020.115369.

30. den Braver-Sewradj, S. P., van Spronsen, R., Hessel, E. V. S., Substitution of bisphenol A: a review of the carcinogenicity, reproductive toxicity, and endocrine disruption potential of alternative substances. Critical Reviews in Toxicology, 2020, 50(2), 128–147. doi: 10.1080/10408444.2019.1701986.

31. Morrissey, R. E., George, J. D., Price, C. J., Tyl, R. W., Marr, M. C., Kimmel, C. A., The developmental toxicity of bisphenol A in rats and mice. Fundamental and Applied Toxicology, 1987, 8(4), 571–582. doi: 10.1016/0272-0590(87)90142-4.

