## Supplementary figures and images for "Bisphenol AF induces overactivation of primordial follicles via Hippo signaling and causes premature ovarian insufficiency in mice"

### Figure S1

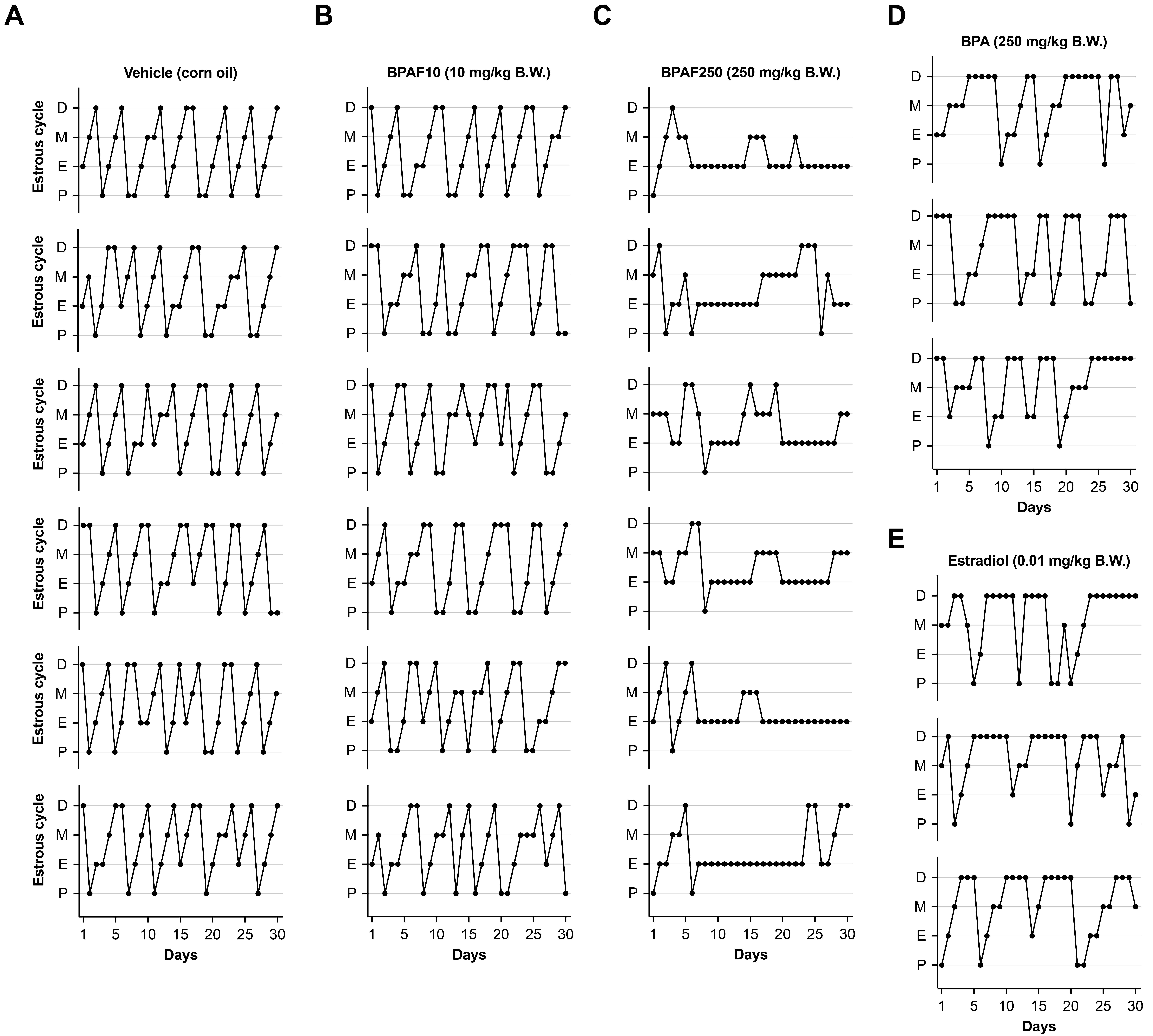

### Figure S2

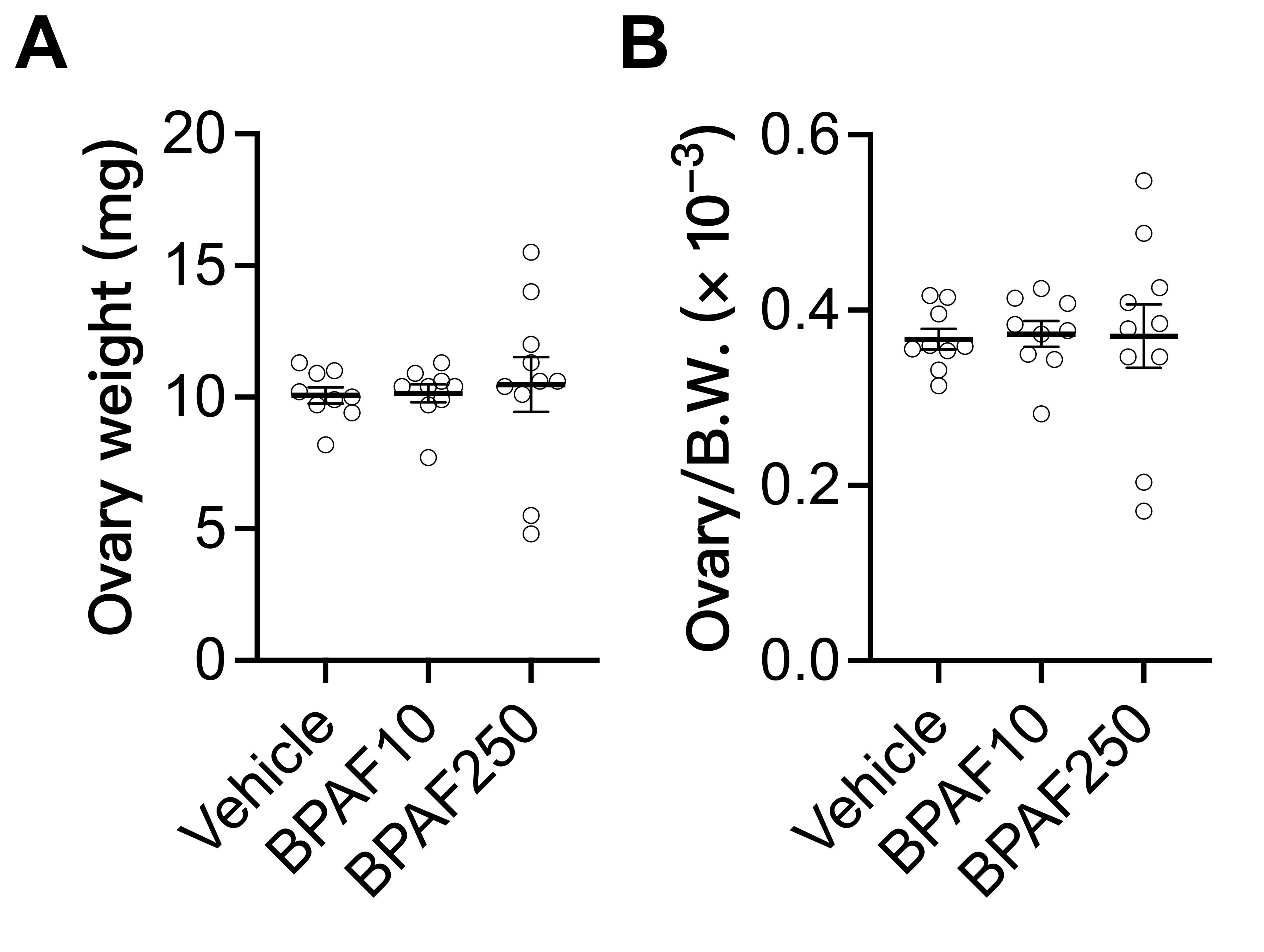

### Figure S3

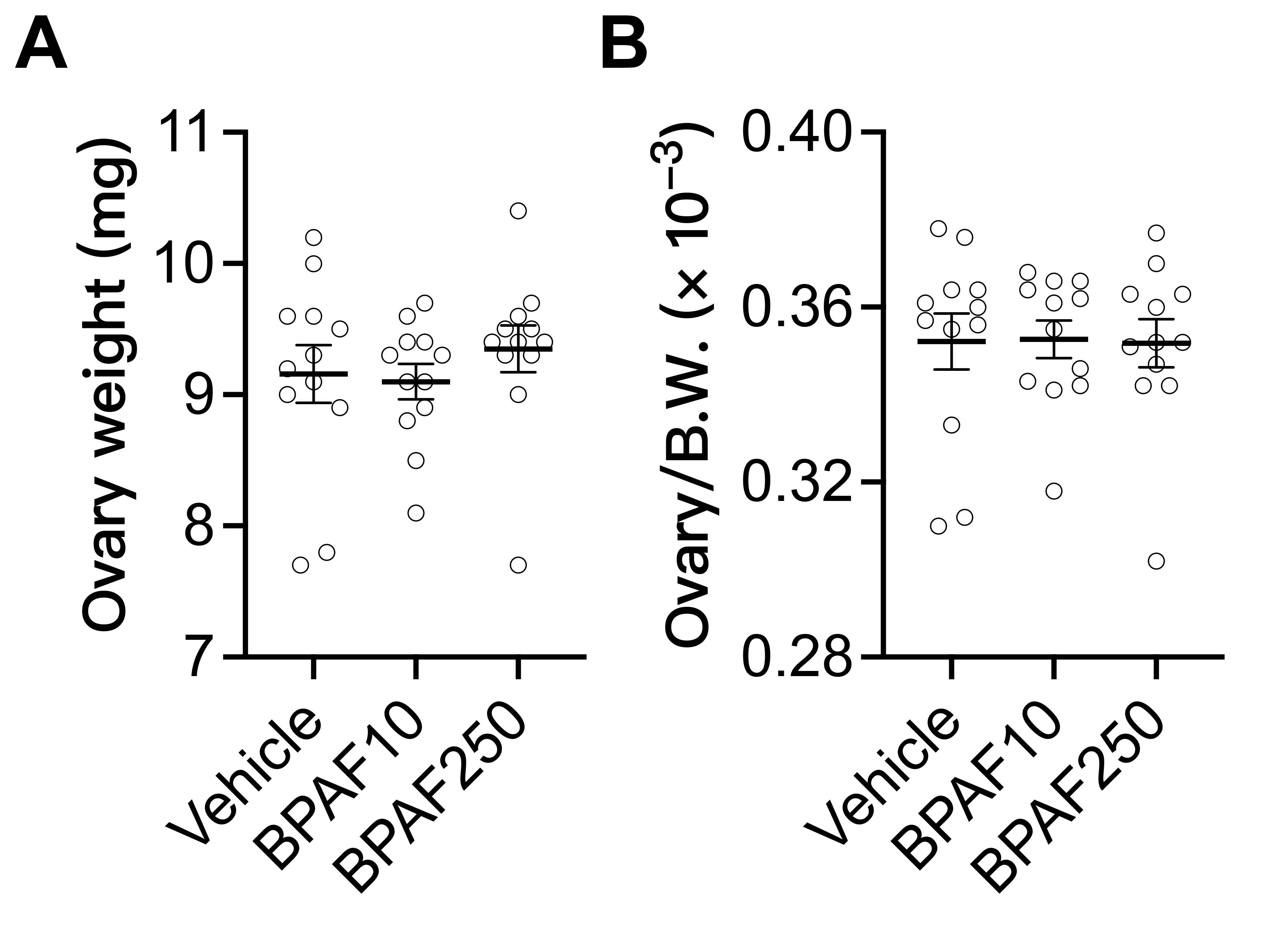
